# *XAP5 CIRCADIAN TIMEKEEPER* regulates RNA splicing and the circadian clock via genetically separable pathways

**DOI:** 10.1101/2022.12.20.521243

**Authors:** Hongtao Zhang (张弘韬), Roderick W. Kumimoto, Shajahan Anver, Stacey L. Harmer

## Abstract

The circadian oscillator allows organisms to synchronize their cellular and physiological activities with diurnal environmental changes. In plants, the circadian clock is primarily composed of multiple transcriptional-translational feedback loops. Regulators of post-transcriptional events, such as pre-mRNA splicing factors, are also involved in controlling the pace of the clock. However, in most cases the underlying mechanisms remain unclear. We have previously identified *XAP5 CIRCADIAN TIMEKEEPER* (*XCT*) as an *Arabidopsis thaliana* circadian clock regulator with uncharacterized molecular functions. Here, we report that XCT physically interacts with components of the spliceosome, including members of the Nineteen Complex (NTC). PacBio Iso-Seq data show that *xct* mutants have transcriptome-wide pre-mRNA splicing defects, predominantly aberrant 3’ splice site selection. Expression of a genomic copy of *XCT* fully rescues those splicing defects, demonstrating that functional *XCT* is important for splicing. Dawn-expressed genes are significantly enriched among those aberrantly spliced in *xct* mutants, suggesting that the splicing activity of *XCT* may be circadian regulated. Furthermore, we show that loss of function mutations in *PRP19A* or *PRP19B*, two homologous core NTC components, suppress the short circadian period phenotype of *xct-2*. However, we do not see rescue of the splicing defects of core clock genes in *prp19 xct* mutants. Therefore, our results suggest that *XCT* may regulate splicing and the clock function through genetically separable pathways.

## Introduction

Most eukaryotes have evolved an endogenous timekeeper known as the circadian clock, which allows them to anticipate the daily fluctuating environmental conditions caused by the earth’s rotation (Harmer, 2009). Although the central oscillators of circadian clocks in diverse eukaryotes lack conserved individual components, they share similar general architectures (Nohales and Kay, 2016). In plants, the approximately 24-h periodicity of the clock is maintained by a complex gene regulatory network consisting primarily of repressors and activators of transcription. Those regulators, often referred to as core circadian clock genes, regulate each other’s expression and the expression of thousands of output genes (Creux and Harmer, 2019).

Additionally, post-transcriptional and post-translational mechanisms such as alternative splicing of precursor messenger RNAs (pre-mRNAs) provide critical regulation of clock function. It has been suggested that changes in splicing of core circadian clock genes are important for Arabidopsis in response to environmental stresses (James et al., 2012; Kwon et al., 2014). For example, cold temperatures significantly suppress production of a splice variant of *CIRCADIAN CLOCK-ASSOCIATED1* (*CCA1*) in which intron four is retained. This incompletely spliced isoform encodes a truncated protein which competitively inhibits the function of fully spliced *CCA1* (Seo et al., 2012). However, in most cases the effects of alternative splicing of pre-mRNAs on circadian clock function remain unclear.

In contrast, the mechanisms underlying pre-mRNA splicing are increasingly well understood (Wilkinson et al., 2020). There are two catalytic transesterification steps, which sequentially remove first the 5’ and then the 3’ splice sites (5’SS and 3’SS) of introns from their adjacent exons (Shi, 2017). This process is carried out by five small nuclear ribonucleoproteins (snRNPs) and hundreds of non-snRNP splicing factor proteins which assemble on a pre-mRNA to form the spliceosome complex (Wilkinson et al., 2020). One of these spliceosome complex components is the Nineteen Complex (NTC, also known as PRP19 complex), named after PRECURSOR RNA PROCESSING 19 (PRP19), a U-box E3 ubiquitin ligase that forms the core of the NTC (Hatakeyama et al., 2001; Koncz et al., 2012). The NTC is associated with the spliceosome during the two transesterification steps and helps facilitate conformational rearrangements and promote splicing fidelity (Fig. 1; Hogg et al., 2010). The NTC is highly conserved across eukaryotes. In Arabidopsis, multiple orthologs of yeast NTC proteins including two PRP19 homologs (PRP19A/MAC3A and PRP19B/MAC3B) have been shown to physically interact with the spliceosome (Monaghan et al., 2009; Deng et al., 2016). Plants mutant for *prp19a prp19b* or homologs of other NTC-associated proteins such as *pleiotropic regulatory locus1* (*prl1*) and *snw/ski-interacting protein* (*skip*) have genome-wide intron retention defects (Jia et al., 2017; Wang et al., 2012; Li et al., 2019). These data indicate that NTC components act as evolutionarily conserved splicing factors in Arabidopsis.

**Figure 1.**
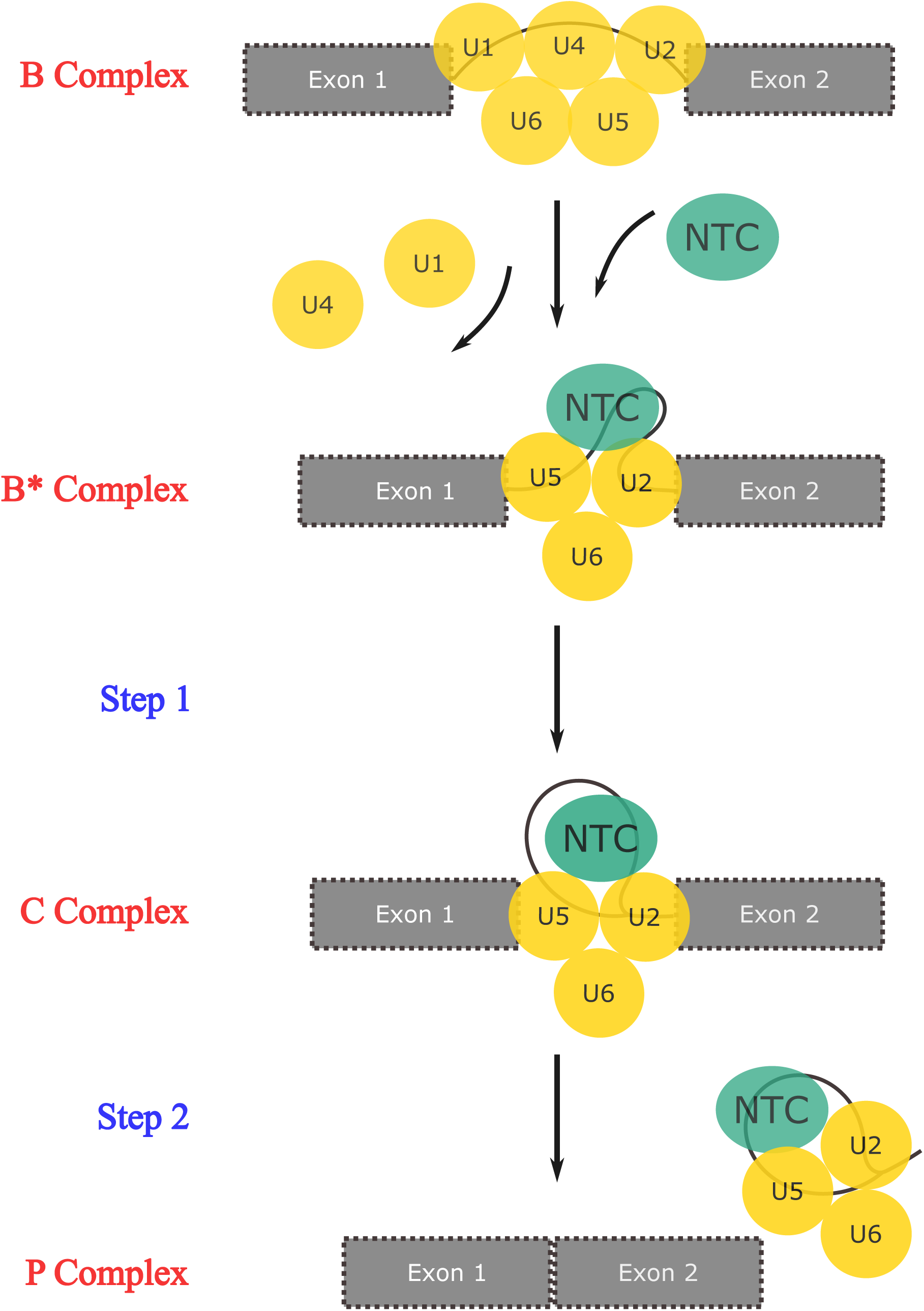
Simplified overview of pre-mRNA splicing reactions. A schematic diagram highlighting the two catalytic transesterification steps and the association between the NTC and the spliceosomal complex during pre-mRNA splicing. Gray boxes and solid black lines represent exons and introns, respectively. U1-U6 small nuclear ribonucleoproteins (snRNPs) are indicated by yellow circles. The NTC is indicated by a green oval.

Mutation of splicing factors can lead to disruption of circadian clock function (Shakhmantsir and Sehgal, 2019). For example, loss-of-function alleles of NTC components, including *PRP19*, *PRL1* and *SKIP*, cause lengthening of circadian period (Wang et al., 2012; Feke et al., 2019; Li et al., 2019). Aberrantly spliced mRNA variants of core circadian clock genes have been detected in these splicing factor mutants (Sanchez et al., 2010; Wang et al., 2012; Jones et al., 2012; Schlaen et al., 2015; Marshall et al., 2016; Li et al., 2019; Feke et al., 2019), suggesting that changes in the pace of the clock might be caused by aberrant splicing of core clock genes. In some cases, epistasis analysis suggests that this may be true (Marshall et al., 2016; Schlaen et al., 2015). However, in other cases, genetic analysis has either not been performed or has revealed additive interactions between the splicing factor and clock gene mutants. In addition, the levels of mis-spliced mRNA variants of clock genes are usually only a small fraction of total transcripts (Jones et al., 2012; Perez-Santángelo et al., 2014; Feke et al., 2019; Sanchez et al., 2010; Wang et al., 2012). Thus, it is unclear whether splicing factors affect clock function solely by controlling the splicing of core clock genes.

RNA splicing factors often function in multiple biological pathways: plants deficient for components of the NTC have defects in immunity, microRNA biogenesis, DNA damage response and transcriptional elongation (Koncz et al., 2012; Monaghan et al., 2009; Zhang et al., 2013; Jia et al., 2017). Some splicing factors are known to carry out roles in nuclear processes biochemically separable from their roles in splicing. In addition to its structural role in the spliceosome, PRP19 also senses DNA damage and promotes DNA repair via its ubiquitin ligase activity (Maréchal et al., 2014). Additionally, SKIP acts in two distinct complexes to regulate splicing and the transcription of genes involved in flowering time control (Li et al., 2019). Nonetheless, whether splicing factors might control circadian clock function via splicing-independent activities has not been investigated.

We previously identified *XAP5 CIRCADIAN TIMEKEEPER* (*XCT*) as a novel regulator of the Arabidopsis circadian clock (Martin-Tryon and Harmer, 2008). Like NTC components, XCT is also well-conserved across eukaryotes. Homologs of XCT (also known as XAP5 proteins) share a highly conserved C-terminal protein domain and are nuclear-localized (Martin-Tryon and Harmer, 2008; Anver et al., 2014; Li et al., 2018; Lee et al., 2020). Previously, we found that transgenic expression of Arabidopsis *XCT* fully rescued the slow-growth phenotype of fission yeast mutant for *xap5* (Anver et al., 2014). Taken together, these data suggest that XCT homologs might share similar molecular and cellular functions across kingdoms.

It has been reported that FAM50A, one of the two XCT orthologs in humans, physically associates with the spliceosomal C complex and its mutants have defects in RNA splicing (Bessonov et al., 2008; Lee et al., 2020). Similarly, a recent study in Arabidopsis also reported the association of XCT with the spliceosome (Liu et al., 2022), implying that XAP5 proteins may be evolutionarily conserved splicing factors. However, evidence suggests that XAP5 proteins may also participate in fundamental biological processes other than splicing. Fission yeast and *Chlamydomonas* XAP5 proteins associate with chromatin and directly regulate transcription (Anver et al., 2014; Li et al., 2018). In addition to its role in the clock, Arabidopsis *XCT* has been implicated in diverse processes including small RNA production, immune signaling, and DNA damage responses (Fang et al., 2015; Xu et al., 2017; Kumimoto et al., 2021). Notably, NTC components have also been reported to function in all these pathways (Jia et al., 2017; Maréchal et al., 2014; Chanarat and Sträßer, 2013). Although pleiotropic defects are seen in *xct* mutants, interconnections between these phenotypes and the molecular function of *XCT* have not been extensively studied.

In the current work, we report that XCT physically interacts with NTC and other spliceosome-associated proteins in Arabidopsis. We use long-read RNA sequencing to reveal that *XCT* controls the fidelity of 3’ splice site selection for hundreds of pre-mRNA splicing events, probably by rejecting downstream suboptimal 3’ splice sites. Intriguingly, circadian-regulated genes that are aberrantly spliced in *xct* mutants are significantly enriched for peak expression at subjective dawn. This implies that the splicing-related activity of *XCT* may be circadian regulated. We also demonstrate that *PRP19A* and *PRP19B* display epistatic genetic interactions with *XCT* in control of the circadian clock period length but not of the splicing of core clock genes. This is consistent with our finding that although *xct-2* causes more severe splicing defects than *xct-1*, *xct-1* mutants have a stronger circadian phenotype. Collectively, our results suggest that XCT works close to NTC and may regulate pre-mRNA splicing and the circadian clock function via distinct pathways.

## Results

### XCT physically interacts with the Nineteen Complex and other spliceosome-associated proteins

To uncover the molecular functions of XCT, we carried out mass spectrometry (MS) experiments to identify its possible interactors. We expressed epitope-tagged XCT protein under the control of the native *XCT* promoter in *xct-2* mutant background (*xct-2 XCT*). This transgene largely rescued the morphological and circadian clock defects of *xct-2* (Supplemental Fig. 1). Next, we affinity purified tagged XCT from plant extracts and analyzed co-purifying proteins via MS. In total, 26 proteins were significantly enriched in the XCT immunoprecipitation compared to control immunoprecipitations (Welch’s two sample *t*-test, *P* < 0.05; Supplemental Dataset 1). Among those XCT-associated proteins, 15 (57.7%) were annotated as mRNA splicing-related, including 11 core NTC or NTC-associated proteins (Table 1), consistent with the MS and co-immunoprecipitation data from a recent report (Liu et al., 2022). Notably, studies in human and yeast have revealed that NTC is physically associated with the spliceosomal complex throughout the two catalytic transesterification steps (Fig. 1; Hogg et al., 2010). Taken together, those results imply that *XCT* may act close to NTC and function in the catalytic steps during pre-mRNA splicing. Interestingly, we also observed 8 (30.8%) chloroplast proteins enriched in the XCT immunoprecipitation (Supplemental Dataset 1), which may be related to the delayed leaf greening phenotypes observed in *xct-2* mutant plants (Martin-Tryon and Harmer, 2008).

**Table 1.**
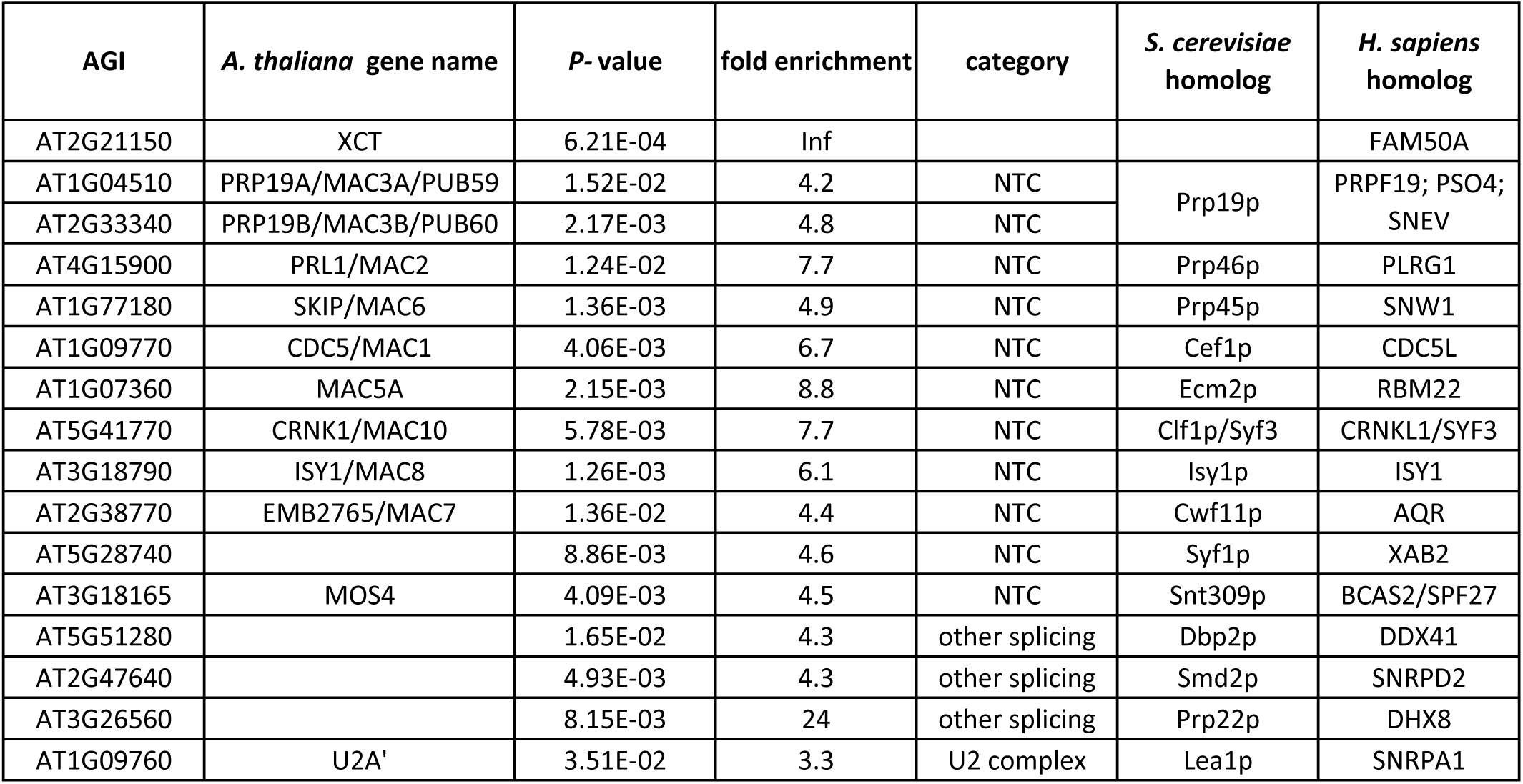
XCT physically associates with splicing-related proteins, especially the NTC components, in Arabidopsis. Mass Spectrometry (MS) data showing that spliceosome-associated proteins are significantly more enriched in XCT-YFP-HA-IP than control IP (fold enrichment > 3; P -value < 0.05, Welch two sample t.test). The full list of proteins co-purified with XCT are described in Supplemental Dataset 1. Only proteins detected in each biological replicate and with 20 or more total peptides counts are shown.

### XCT is required for the fidelity of 3’ splice site selection during pre-mRNA splicing

To investigate a possible role for *XCT* in RNA splicing, we performed PacBio Isoform Sequencing (Iso-Seq) on wild-type Col-0, reduction-of-function allele *xct-1*, null allele *xct-2*, and the complemented line *xct-2 XCT*. Additionally, we also sequenced *prl1-2*, which contains a T-DNA insertion mutation *in PRL1*, an NTC member that has been demonstrated to control genome-wide splicing efficiency (Jia et al., 2017). Since both transcript levels and RNA splicing of a large proportion of the Arabidopsis transcriptome are circadian regulated (Romanowski et al., 2020; Yang et al., 2020), the time of day at which plants are harvested has significant effects on gene expression and splicing analysis (Hsu and Harmer, 2012). Therefore, we grew plants in constant environmental conditions for ten days to desynchronize clocks of individual plants. To further minimize any differences in subjective time of day between wild-type plants, the long-circadian-period *prl1-2*, and the short-period *xct* mutants (Supplemental Fig. 2), we pooled seedlings harvested at 2-hour intervals across a twenty-four hour period (Fig. 2A). For each genotype, we sequenced three biological replicates and acquired an average of 784,832 full-length transcript reads per genotype per replicate (Supplemental Dataset 2). With no alignment needed, those reads were directly mapped to the TAIR 10 Arabidopsis reference genome and then subjected to differential splicing analysis using the JunctionSeq R package (Hartley and Mullikin, 2016).

**Figure 2.**
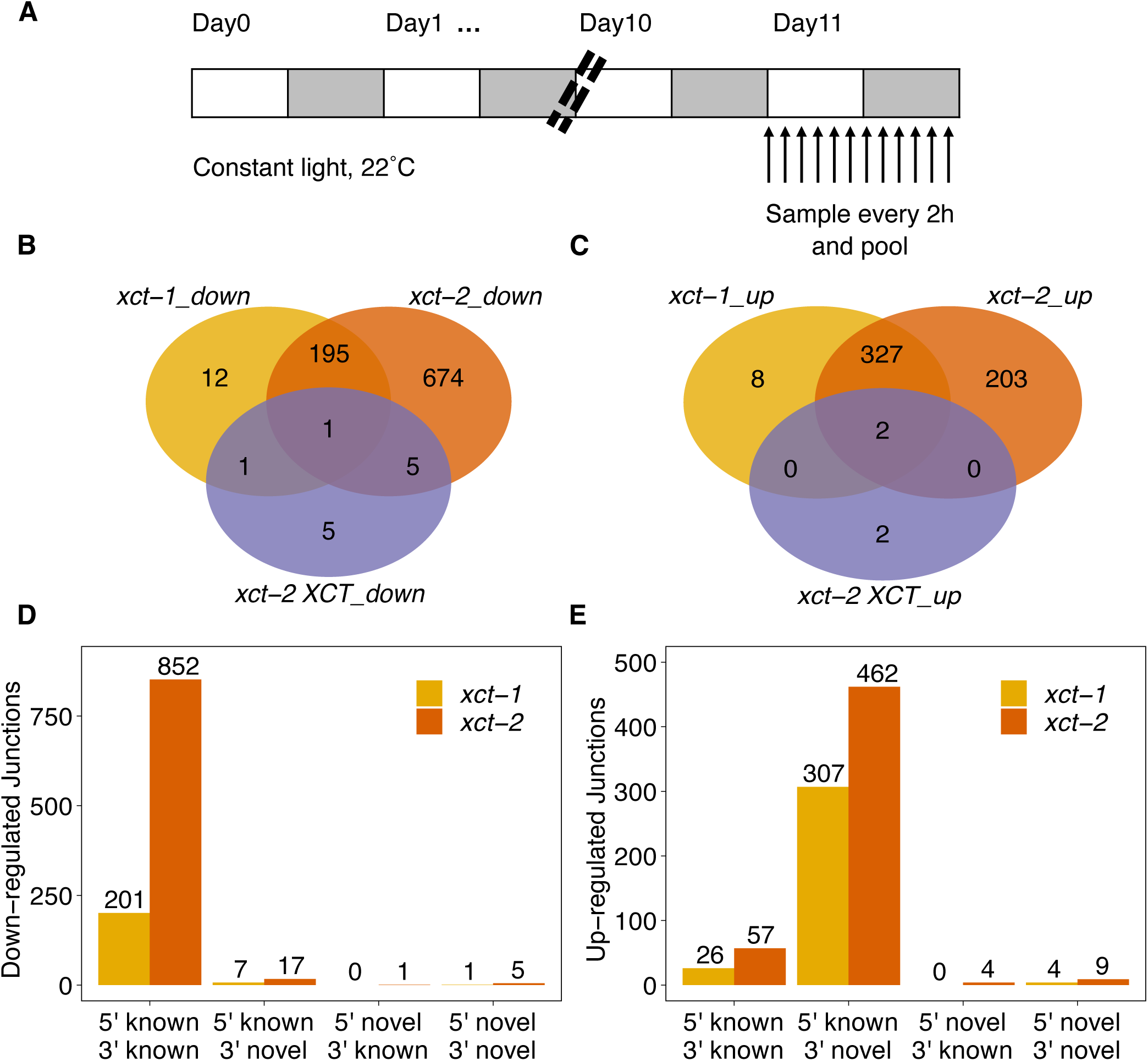
Transcriptome-wide analysis reveals *XCT* as a global pre-mRNA splicing regulator. A, Experimental design and sampling method for the PacBio Iso-Seq experiment. White and grey boxes represent subjective day and subjective night, respectively. Arabidopsis seedlings were grown at constant light and temperature for 10 days before being harvested and pooled. B and C, Numbers of differentially enriched splicing events represented by significantly differentially down-(B) or up-regulated (C) splice junctions in *xct-1*, *xct-2* and *xct-2* complemented with p*XCT*::g*XCT-YFP-HA* compared with Col-0 (false discovery rate < 0.05). D and E, Frequency of different classes of 5’ and 3’ splice sites among down-(D) or up-regulated (E) splice junctions in *xct-1* and *xct-2*. Known or novel splice sites were classified by comparing to TAIR10 genome annotation.

Previous studies have demonstrated that intron retention is the most prevalent type of alternative splicing event in Arabidopsis (Wang and Brendel, 2006; Filichkin et al., 2010). Indeed, plants mutant for multiple NTC components have been reported to have global intron retention defects (Jia et al., 2017; Meng et al., 2022). In our analysis of *prl1-2*, we found 5,730 out of 44,496 detected splicing events were significantly differentially enriched from Col-0 (Supplemental Dataset 3). Notably, 99% of these events were decreases in known junctions (Supplemental Fig. 3), consistent with previous studies suggesting that loss of *PRL1* function mainly causes intron retention without affecting splice site selection (Jia et al., 2017).

**Figure 3.**
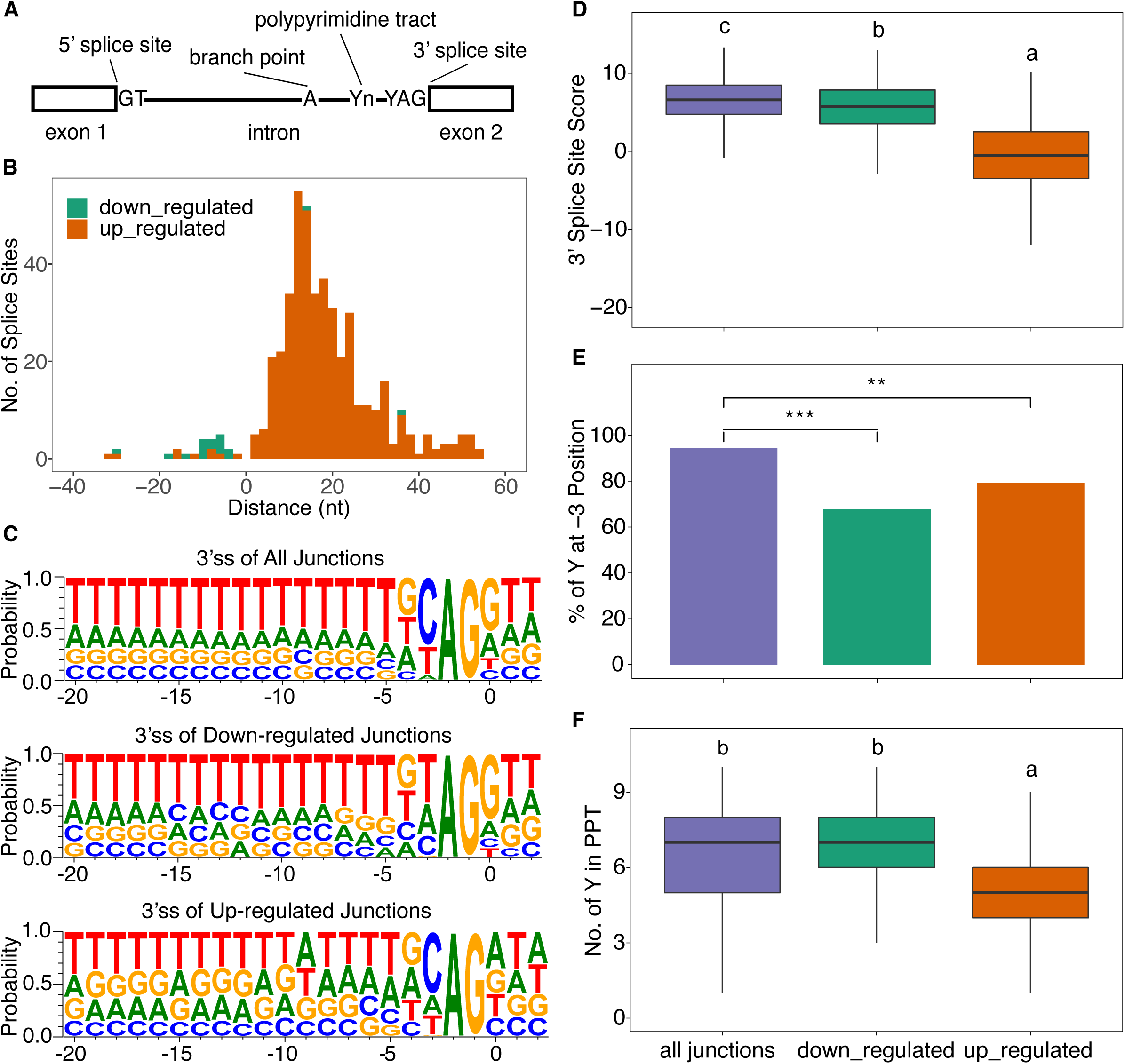
*XCT* is required for the fidelity of 3’ splice site selection during pre-mRNA splicing. A, A schematic diagram showing the structure of a typical U2-type splice junction. Y: pyrimidine. B, Distribution of the distance between each pair of novel 3’ splice site and its corresponding canonical 3’ splice sites in *xct-2*. C, Pictograms showing the frequency of nucleotides in the 23-mers sequences flanking the 3’ splice sites in all expressed, down-regulated and up-regulated splice junctions in *xct-2*. D, Maximum Entropy Model scores showing the strength of 3’ splice sites of all expressed, down-regulated and up-regulated junctions in *xct-2*. E, Percentage of pyrimidines at the -3 position (i.e. the nucleotide preceding the AG at 3’ splice site) in *xct-2*. F, Counts of pyrimidines in the Y10 polypyrimidine tract upstream of the 3’ splice sites in *xct-2*. PPT, polypyrimidine tract. The lines in the boxplot represent the 75% quartile, median and 25% quartile of the data, respectively. Statistical significance in (D) and (F) was determined using linear regression model with junction class as a fixed effect and is shown in lower case letters (Tukey’s multiple comparison test, *P* < 0.05). Statistical significance in (E) was determined by Fisher’s exact test: **: *P* < 0.01; ***: *P* < 0.001.

We next examined transcript composition in *xct* mutants. In the null allele *xct-2*, we detected 75,073 splicing events, of which 875 were significantly down-regulated and 532 were up-regulated compared with Col-0 (FDR<0.05, Fig. 2B-C). Meanwhile, 209 and 337 splicing events were significantly decreased or increased, respectively, in the partial loss-of-function mutant *xct-1*. Comparing mis-regulated splicing between the two *xct* mutant alleles, 196 (93.8%) of the down-regulated and 329 (97.6%) of the up-regulated events were shared. Thus, the two *xct* alleles have similar splicing phenotypes but the severity of the defect is greater in the null allele despite the stronger effect of *xct-1* on circadian period (Supplemental Fig. 1). Next, we analyzed the splicing events in the complemented line *xct-2 XCT*. Only 16 out of 84,965 events were significantly differentially enriched compared with Col-0, indicating that the splicing defects of *xct-2* were almost completely rescued by the restored *XCT* expression. Taken together, these results demonstrate that *XCT* is important for transcriptome-wide pre-mRNA splicing.

To further characterize the major types of splicing defects caused by loss of *XCT* function, we analyzed the splice sites for the differentially spliced junctions in *xct* mutants. Specifically, we categorized *xct*-induced mis-splicing events into four classes based on whether the 5’ and 3’ splice sites used were previously annotated (known) or not (novel). As expected, most down-regulated splicing events displayed decreases in usage of junctions with known 5’ and 3’ splice sites (Fig. 2D). Interestingly, 307 (90.8%) in *xct-1* and 462 (86.5%) in *xct-2* of up-regulated splicing events involved increased usage of junctions with a known 5’ splice site but a novel 3’ splice site (Fig. 2E). The decreases in abundance of junctions with known splice sites hence reflects both intron retention events and novel 3’ splice sites usage. Therefore, our results demonstrate that in addition to controlling the efficient removal of introns, *XCT* is also responsible for the fidelity of 3’ splice site selection.

To investigate how *XCT* contributes to the fidelity of 3’ splice site selection, we compared the significantly up-or down-regulated 3’ splice sites in *xct* mutants with all detected 3’ splice sites in Col-0. Intriguingly, most of the up-regulated novel 3’ splice sites in *xct-1* and *xct-2* were located less than 50 nucleotides downstream of the wild-type 3’ splice sites (Fig. 3B; Supplemental Fig. 4A). Studies in human and yeast have demonstrated that the sequences preceding a 3’ splice site, including the polypyrimidine tract (PPT) and the pyrimidine at the -3 position (Fig. 3A), are important for the strength of the 3’ splice site (Horowitz, 2012). Therefore, we further examined the 23-mer sequences flanking the 3’ splice sites (from -20 to +3 position).

**Figure 4.**
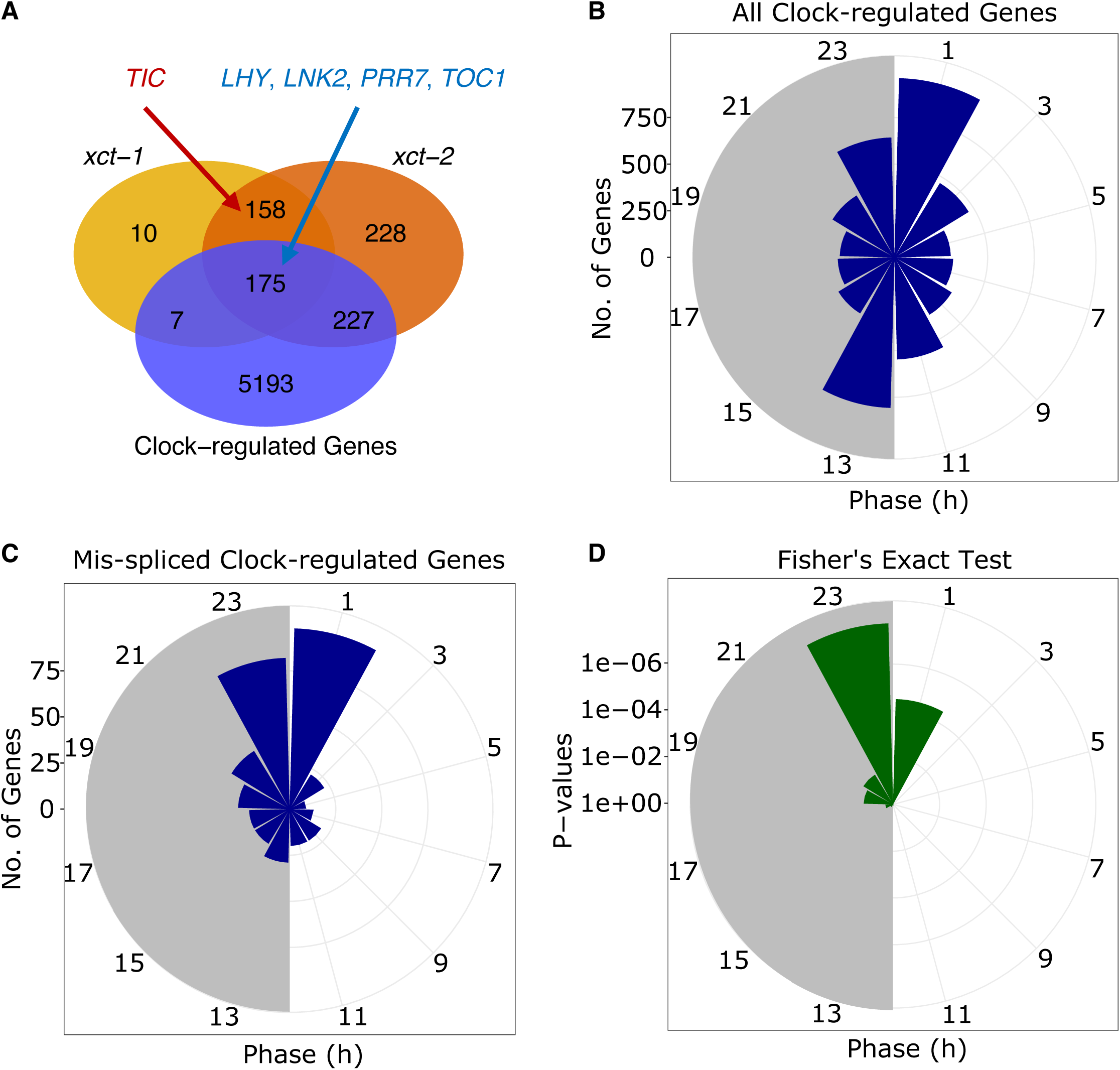
Dawn-phased genes are significantly enriched among all the circadian-clock-regulated genes that are aberrantly spliced in *xct-2.* A, Venn diagram showing the overlaps of aberrantly spliced genes in *xct-1, xct-2*, and total circadian-clock-regulated genes (Romanowski et al., 2020). The core circadian clock genes that are mis-spliced in both *xct* mutants are indicated in red (non-clock-regulated) and blue (clock-regulated) fonts. Only genes considered as ‘detected’ in all three RNA-Seq datasets are included. B and C, Phases of estimated peak expression of all circadian clock-regulated genes (Romanowski et al., 2020) that are detected (B) or significantly aberrantly spliced (C) in *xct-2*. The white and gray backgrounds represent the subjective day and subjective night, respectively. D, Distribution of *P*-values from Fisher’s extact tests calculating whether the ratio of the number of aberrantly spliced genes in (B) to total clock-regulated genes in (A) is significantly higher than expected by chance in each 2-hour interval.

We found that the frequency of the canonical AG sequence at the 3’ splice sites was not altered in either *xct-1* or *xct-2* (Fig. 3C; Supplemental Fig. 4B-C). However, the average predicted strength of the 875 down-regulated 3’ splice sites in *xct-2* was significantly weaker than the average of total detected 3’ splice sites in Col-0 (Fig. 3D). Specifically, the percentage of pyrimidine residues at the -3 position was significantly reduced in down-regulated 3’ splice sites compared to that found in all detected 3’ splice sites (Fisher’s exact test, *P* < 0.001, Fig. 3E), suggesting that functional *XCT* is important for the removal of 3’ splice sites with a suboptimal sequence at the -3 position. Moreover, the 532 up-regulated 3’ splice sites had an even lower 3’ splice site score than the down-regulated sites (Fig. 3D). The frequency of pyrimidines both at the -3 position and throughout the PPT region was significantly lower in the up-regulated junctions (Fig. 3E-F), showing that *xct-2* is less able to discriminate between 3’ splice sites during splicing. Taken together, our Iso-Seq data demonstrates that *XCT* controls the accuracy of 3’ splice site selection, possibly by helping to recognize weaker 3’ splice sites and reject sub-optimal downstream sites.

### The splicing defects of core clock genes are generally more severe in xct-2 than xct-1

Since aberrant splicing of core clock genes may fully or partially contribute to the altered circadian clock period phenotype in splicing factor mutants (Sanchez et al., 2010; Wang et al., 2012; Jones et al., 2012; Schlaen et al., 2015; Marshall et al., 2016; Li et al., 2019; Feke et al., 2019), we searched for core clock genes with aberrant splicing events in both the short-period *xct-1* and *xct-2* mutants. In total, our Iso-Seq data detected five aberrantly spliced core clock genes, including *LHY*, *LNK2*, *PRR7*, *TOC1* and *TIC* (Fig. 4A). However, we noticed that the ratio of aberrantly-spliced relative to fully-spliced transcripts for several clock genes are significantly higher in *xct-2* than *xct-1*, even though the circadian period defect is more severe in the latter (Supplemental Fig. 5A-D; Supplemental Fig. 1B). We confirmed this finding via semi-qRT-PCR assay (Supplemental Fig. 5E-H). Thus, those results imply that there may be a disconnect between the splicing and circadian clock defects in *xct-1* and *xct-2*.

**Figure 5.**
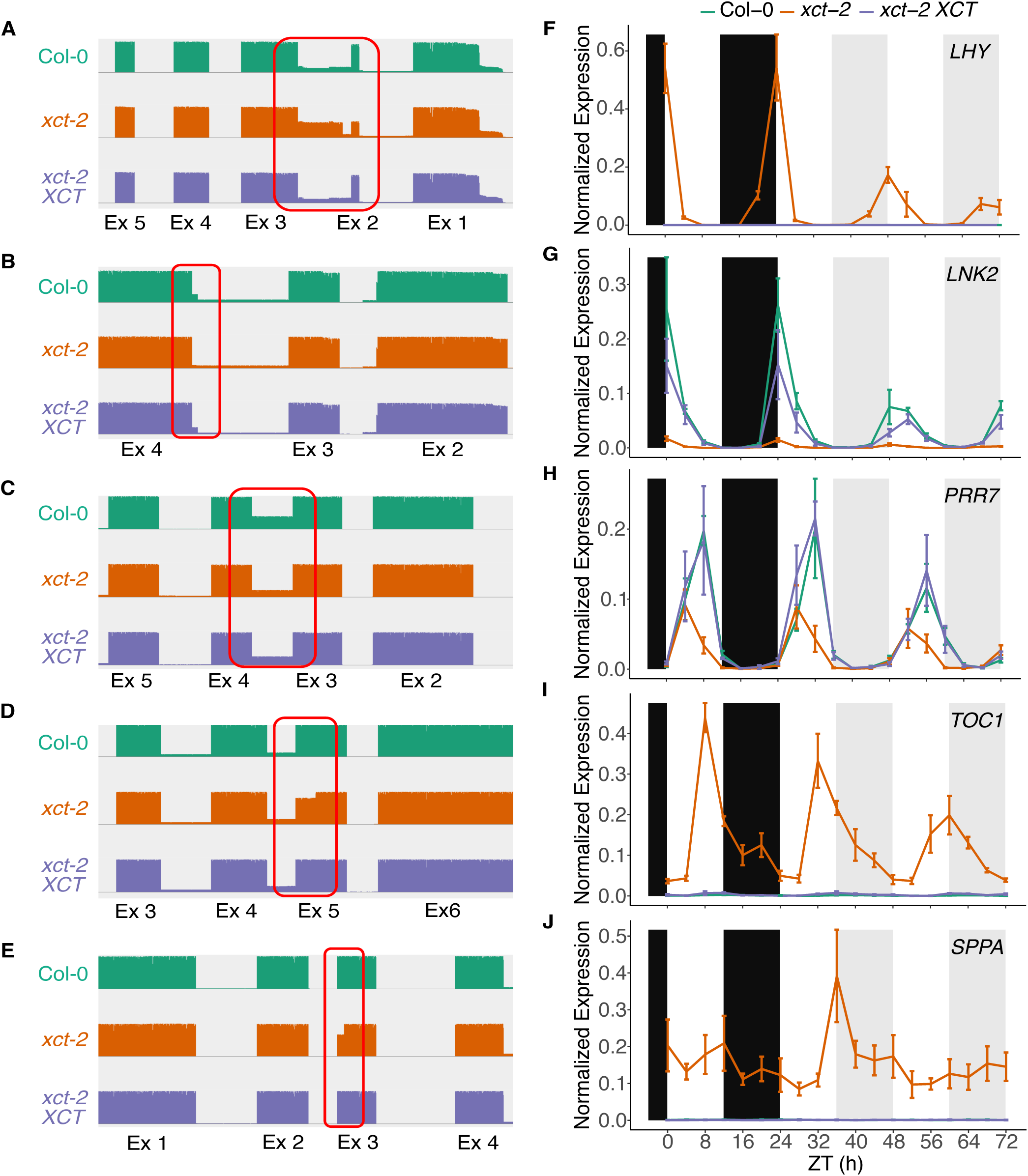
Time-course qRT-PCR experiments validate the role of *XCT* in regulating pre-mRNA splicing of clock-regulated and non-clock-regulated genes. A -E, Sashimi plots showing PacBio Iso-Seq reads mapped to *LHY* (A)*, LNK2* (B)*, PRR7* (C)*, TOC1* (D) and *SPPA* (E) in Col-0 (teal)*, xct-2* (orange) and *xct-2 XCT* (purple). The red rectangles highlight the aberrantly spliced exon-exon junctions that are examined by qRT-PCR in (F) - (J). F - J, Normalized expression of the aberrantly spliced isoforms of *LHY* (F)*, LNK2* (G)*, PRR7* (H)*, TOC1* (I) and *SPPA* (J) in Col-0*, xct-2* and *xct-2 XCT*. Samples were collected every four hours over a 72-h window. Expression levels were examined by qRT-PCR using splice-junction- specific primers and normalized to *PP2*A and *IPP2.* Data points represent mean ± se from three independent biological replicates. For each isoform in each biological replicate, the normalized expression levels were relative to the highest expression levels of their total transcripts in Col-0 across all time points. Teal lines, wild type Col-0; orange lines, *xct-2* mutants; purple lines, *xct-2 XCT*. Black background, dark period; white and gray background, light period during subjective day and night, respectively.

### Genes expressed at subjective dawn are enriched among those aberrantly spliced in xct mutants

To more broadly explore possible links between the circadian clock and splicing phenotypes in *xct* mutants, we looked for enrichment of circadian clock regulation within genes aberrantly spliced in *xct.* Among the 5,602 previously reported clock-regulated genes (Romanowski et al., 2020) that are detected in our Iso-Seq data, we identified 182 and 402 genes that are mis-spliced in *xct-1* and *xct-2*, respectively (Fig. 4A; Supplemental Dataset 4). However, there is no significant over-representation in either mutant (one-tailed Fisher’s exact test, *P* = 0.87 for *xct-1* and 0.99 for *xct-2*), suggesting that *XCT* does not preferentially affect the splicing of clock-regulated genes.

Previous studies have identified transcripts that are differentially spliced at various times of day (Romanowski et al., 2020; Yang et al., 2020). This suggests that splicing activity may be circadian clock regulated. Therefore, to test whether *XCT* preferentially affects pre-mRNA splicing at certain times of day, we examined the distribution of estimated peak expression times of genes aberrantly spliced in *xct* mutants. We grouped all 5,602 clock-regulated genes into twelve 2-hour intervals based on their estimated peak expression time (Fig. 4B). Genes that are aberrantly spliced in *xct-2* are significantly enriched for peak expression between ZT22 and ZT2 compared with the other intervals of the day (Fig. 4C-D, Fisher’s exact test, *P* < 1e-4). Similarly, clock-regulated genes that are aberrantly spliced in *xct-1* also showed a dawn-enriched expression pattern (Supplemental Fig. 6). Collectively, those results indicate that *XCT* activity within the spliceosome may be regulated by the circadian clock.

**Figure 6.**
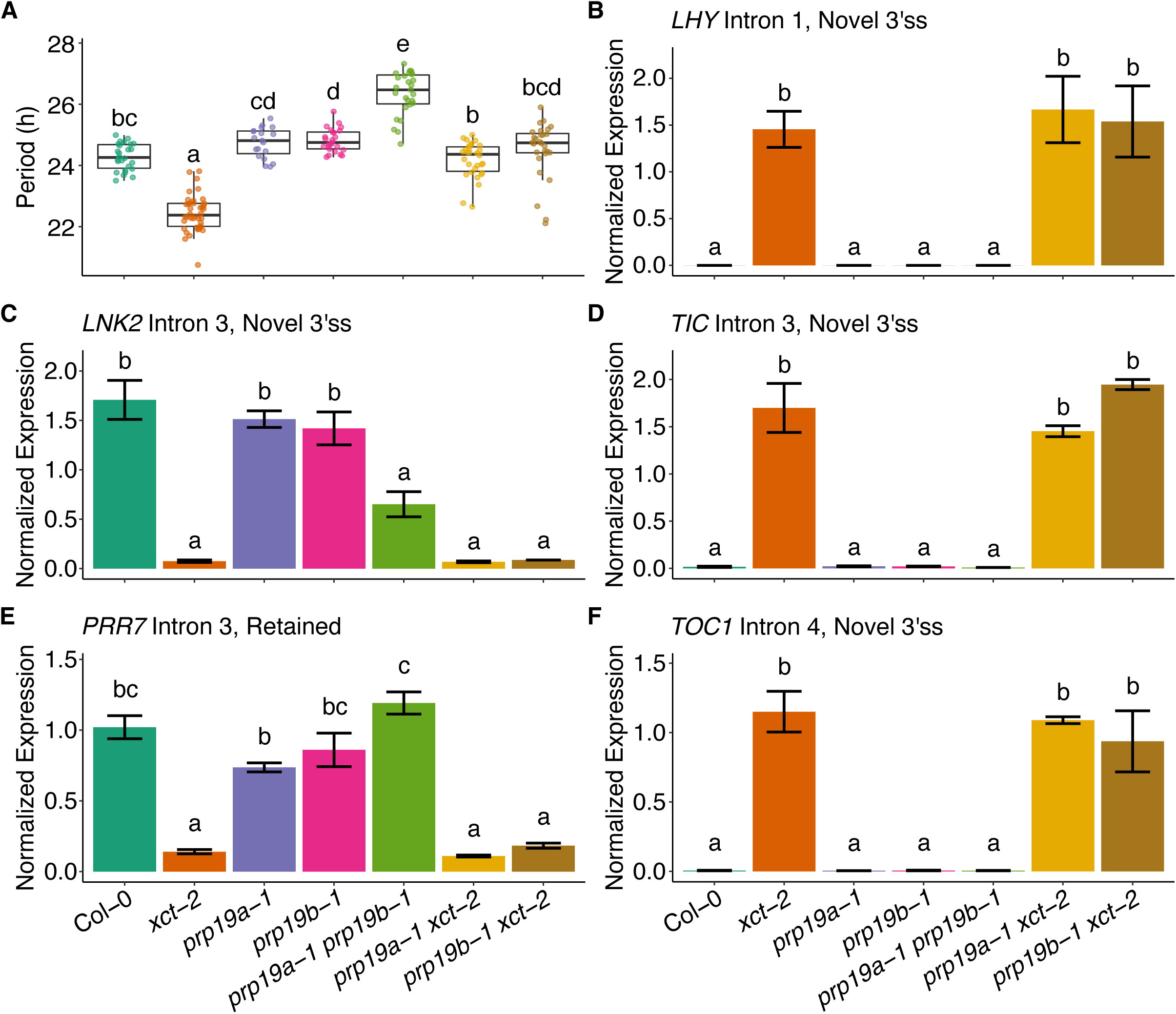
Loss of *PRP19* function suppresses the short circadian clock period phenotype but not the splicing defects of core clock genes in *xct-2*. A, Circadian periods of Col-0, *xct-2*, *prp19a-1*, *prp19b-1*, *prp19a-1 prp19b-1*, *prp19a-1 xct-2* and *prp19b-1 xct-2* plants. B - F, Normalized expression of the aberrantly spliced isoforms of *LHY, LNK2, TIC, PRR7* and *TOC1* in Col-0, *xct-2*, *prp19a-1*, *prp19b-1*, *prp19a-1 prp19b-1*, *prp19a-1 xct-2* and *prp19b-1 xct-2*. Samples were collected at the estimated peak expression time for each gene. Expression levels were examined by qRT-PCR using splice-junction-specific primers and normalized to *PP2*A and *IPP2.* Data points represent mean ± se from two independent biological replicates. Statistical significance was determined using linear regression model with genotype as a fixed effect and is shown in lower case letters (Tukey’s multiple comparison test, *P* < 0.05).

To further investigate whether *XCT* preferentially affects splicing of transcripts produced at dawn, we monitored the abundance of candidate splice variants via quantitative reverse transcriptase-polymerase chain reaction (qRT-PCR) over a 24-hour light/dark period (LD) followed by 48 hours in constant light (LL). We selected four circadian-clock-regulated genes and one non-clock-regulated gene that are aberrantly spliced in *xct-2* and fully rescued in *xct-2 XCT* (Fig. 5A-E). Total transcript levels of clock-regulated genes displayed rhythmic abundance with peak *LHY* and *LNK2* expression at dawn and peak *PRR7* and *TOC1* expression late in the afternoon (Supplemental Fig. 7), as expected (Creux and Harmer, 2019). We found significantly advanced phases of expression in *xct-2* but this mutation did not affect the overall abundance of total transcripts for those genes. However, both the peak and average abundance of the aberrantly spliced isoforms were significantly different in *xct-2* compared with Col-0 (Fig. 5F-J). This serves as validation of the pooling strategy we used to generate our Iso-Seq libraries, where the daily abundance of transcripts was averaged (Fig. 2A; Fig. 5A-E). Contrary to our expectations, instead of preferentially accumulating at dawn, abundance of all splice variants in *xct-2* was synchronized with total transcript levels (Fig. 5F-I; Supplemental Fig. 7). Likewise, the mis-spliced isoform of the non-clock-regulated gene *SPPA* did not show a significant morning peak (Fig. 5J). Meanwhile, the genomic *XCT* complemented line almost completely rescued the splicing defects across the whole experimental period. Taken together, those results suggest that although morning-expressed genes are preferentially enriched among *XCT* targets, *XCT* is important for the accuracy of pre-mRNA splicing throughout the day under both LD and free-running LL conditions.

### Reduction of PRP19 function rescues the circadian clock but not the splicing defects in xct-2

Previous studies revealed that NTC components participate in regulation of circadian clock function in Arabidopsis (Feke et al., 2019; Li et al., 2019). Here we examined circadian period in mutants for several NTC components. We found that mutation in either of the two Arabidopsis *PRP19* homologs, *PRP19A* (*MAC3A*) or *PRP19B* (*MAC3B*), only caused minor lengthening of circadian period. In contrast, the *prp19a-1 prp19b-1* double mutant had a significantly slower circadian clock than Col-0 (Supplemental Fig. 8A), consistent with previously reported redundant roles for *PRP19A* and *PRP19B* in circadian clock regulation (Feke et al., 2019). Similarly, loss of function of other NTC members, including *PRL1*, *CDC5* and *SKIP*, also lengthened the clock period by 1 to 3 hours (Supplemental Fig. 2B; Supplemental Fig. 8B-C), similar to previous reports for *prl1-9* and *skip-1* mutants (Li et al., 2019).

Since *PRP19* co-purifies with *XCT* (Table 1) and they both control the pace of the circadian clock (Supplemental Fig. 8A), we hypothesized they might function in the same pathway to regulate the clock. To test this hypothesis, we introduced *prp19a-1* and *prp19b-1* mutant alleles, which express greatly decreased and near-null levels of *PRP19A* and *PRP19B*, respectively (Supplemental Fig. 9), into the null *xct-2* background and assayed their circadian clock phenotypes. Interestingly, we found that neither the circadian period of *prp19a-1 xct-2* nor of *prp19b-1 xct-2* was significantly different from Col-0 (Fig. 6A; Supplemental Fig. 10), indicating that both *prp19* mutants can fully suppress the short-period phenotype of *xct-2*. Notably, *prp19a-1* and *prp19b-1* single mutants only have slightly longer circadian periods than Col-0. This suggests that the rescued clock function in *prp19 xct* double mutants is not likely due to additive genetic interactions. Thus, our data show that functional *PRP19A* and *PRP19B* are both necessary for *XCT* in regulation of circadian period. To further investigate the functional association between *XCT* and *PRP19*, we next asked whether *prp19* mutants suppress other developmental and physiological defects observed in *xct-2*. In contrast to the circadian period, the reduced rosette size in *xct-2* was not significantly restored in either *prp19a-1 xct-2* or *prp19b-1 xct-2* (Supplemental Fig. 11), indicating that the effects of *XCT* on rosette development and the circadian clock period are genetically separable. This is consistent with our observation that *xct-1* and *xct-2* both have short circadian period phenotypes but only *xct-2* is morphologically different from Col-0 (Supplemental Fig. 1).

Next, to determine whether suppression of the short clock period phenotype in *xct-2 prp19* mutants is due to reversal of the *xct-2* splicing defects, we conducted qRT-PCR experiments to examine the abundance of *xct*-induced aberrantly-spliced mRNA isoforms of core clock genes in *prp19a-1 xct-2* and *prp19b-1 xct-2*. Surprisingly, none of the aberrantly-spliced isoforms tested showed significantly different abundance in *prp19a-1 xct-2* and *prp19b-1 xct-2* relative to *xct-2* (Fig. 6B-F). Consistent with previous reports for functional redundancy between *PRP19A* and *PRP19B* (Monaghan et al., 2009; Li et al., 2019), we noticed a statistically significant defect in splicing of the third intron of *LNK2* in the *prp19a-1 prp19b-1* double mutant but not the *prp19a-1* or *prp19b-1* single mutants. We also checked the expression levels of fully-spliced or total mRNA isoforms of the five core clock genes that are aberrantly-spliced in *xct-2*. In all cases, addition of *prp19a-1* or *prp19b-1* to *xct-2* did not alter the abundance of functionally spliced isoforms of those clock genes (Supplemental Fig. 12). Thus, similar to the developmental defects, the splicing defects of core clock genes are genetically separable from the circadian clock phenotype of *xct-2*.

## Discussion

The accurate removal of introns from pre-mRNAs is an essential step of gene expression in all eukaryotes. In this work, we report that *XCT* is a global regulator of pre-mRNA splicing in Arabidopsis. Multiple lines of evidence suggest that orthologs of XCT share conserved molecular and cellular functions across kingdoms. Previous mass spectrometry data showed that FAM50A and FAM50B, two human homologs of XCT, physically associate with the affinity-purified spliceosomal C complex (Bessonov et al., 2008; Bessonov et al., 2010). Loss of *FAM50A* function induces transcriptome-wide pre-mRNA splicing defects in both human and zebrafish (Lee et al., 2020). Similarly, a recent study in Arabidopsis also found that XCT associates with spliceosomal proteins and regulates splicing (Liu et al., 2022). In this paper, we show that XCT physically co-purifies with PRP19 and other NTC-associated proteins (Table 1). Furthermore, our PacBio Iso-Seq data indicate that the efficiency and fidelity of pre-mRNA splicing are negatively impacted in both the partial loss-of-function mutant *xct-1* and the null mutant *xct-2* but are largely restored in *xct-2 XCT* (Fig. 2). Therefore, XCT orthologs likely play evolutionarily conserved roles in pre-mRNA splicing. It would be interesting to apply structural biology approaches to further investigate what conserved functional domains among XCT homologs might contribute to their splicing activities.

Studies have shown that intron retention is the most prevalent type of alternative splicing event in Arabidopsis (Wang and Brendel, 2006; Filichkin et al., 2010). Correspondingly, mutations in many Arabidopsis splicing factors mainly cause intron retention (Supplemental Fig. 3; Schlaen et al., 2015; Li et al., 2019). Here we identify *XCT* as an unusual splicing regulator that specifically controls the fidelity of 3’ splice site selection (Fig. 2D-E; Fig. 3). Biochemical studies of splicing in yeast revealed that several DEAH-box ATPases are important for the selection of 3’ splice sites (Horowitz, 2012). For example, PRP22, a DEAH-box ATPases that co-purifies with XCT (Table 1; Liu et al., 2022), represses usage of aberrant 3’ splice sites and promotes spliceosome scanning for downstream alternative 3’ splice sites in yeast (Mayas et al., 2006; Semlow et al., 2016). Studies in human showed that hPRP22, FAM50A and FAM50B, are all abundant in the spliceosomal C complex but nearly absent in B and B* complexes (Bessonov et al., 2008; Bessonov et al., 2010; Zhan et al., 2022). This implies that PRP22 and XCT may have evolutionarily conserved interactions during the second transesterification reaction of splicing, when 5’ and 3’ splice sites are joined (Fig. 1). Intriguingly, we detected decreased fidelity of 3’ splice site selection in both *xct-1* and *xct-2* mutants (Fig. 3; Supplementary Fig. 4), which is also observed in yeast *prp22* mutants (Semlow et al., 2016). Therefore, our results imply that XCT may work with PRP22 to control the fidelity of 3’ splice site selection. Future research on the biochemical functions of XCT using *in vitro* and *in vivo* systems could help reveal how it controls splicing fidelity.

Previous microarray and RNAseq studies demonstrated that transcription and splicing of a large proportion of Arabidopsis genes are under circadian clock regulation (Covington and Harmer, 2007; Hsu and Harmer, 2012; Romanowski et al., 2020; Yang et al., 2020). Consequently, timing of sample collection is an important consideration in gene expression and splicing analysis, especially when comparing genotypes with different circadian periods. Here we employed a more efficient sampling method by harvesting plants at twelve evenly distributed time points across a day and pooling them before analysis. Our pooling strategy enabled us to detect transcripts of genes with the distribution of peak phases of expression mirroring that of all clock-regulated genes (Fig. 4B; Romanowski et al., 2020). Overall, this strategy allowed us to identify more clock-regulated differentially spliced transcripts in *xct-2* mutants than in a recent study in which plants were collected at a single time point (Liu et al., 2022). For example, we report novel differential splicing events in afternoon-expressed genes including *PRR7* and *TOC1* (Fig. 4A; Supplemental Fig. 7C-D). In fact, our time-course qPCR data suggest that abundance of the aberrantly spliced isoforms of these clock-regulated genes fluctuates by over 99% depending on time of sample collection (Fig. 5F-I). Thus, our results demonstrate the advantages of pooling samples when conducting transcriptome-wide splicing analysis of clock-regulated genes.

Using this strategy, we were able to investigate whether *XCT* preferentially affects splicing at certain times of day. Indeed, we found that dawn-expressed genes are significantly more likely to be mis-spliced in *xct* mutants than those with other peak phases (Fig. 4C-D; Supplemental Fig. 6). This suggests that *XCT* activity in or its association with the spliceosome may be clock-regulated. An alternative explanation for the overrepresented mis-splicing of dawn-expressed genes could be that *cis*-regulatory elements of those genes might preferentially recruit XCT to control their splicing. This possibility is supported by previous studies showing that XCT orthologs in fission yeast and *Chlamydomonas* are chromatin-associated and in the latter case recruit RNA polymerase II (Pol II) to promoter regions (Anver et al., 2014; Li et al., 2018).

Alternative splicing regulates various biological processes, including the function of the circadian clock (Hsu and Harmer, 2014; Nolte and Staiger, 2015). In Arabidopsis, such regulation is supported by identification of splicing factor mutants that alter circadian clock period length (Shakhmantsir and Sehgal, 2019). Some splicing factors, such as *GEMIN2* and *SICKLE*, interact epistatically with one or more alternatively spliced clock genes in the control of period length (Schlaen et al., 2015; Marshall et al., 2016), suggesting changes in the pace of the clock are due to altered splicing of these clock genes. Whereas in other cases, the mis-regulated circadian period can only be partially attributed to changes in levels of splicing variants of clock genes (Sanchez et al., 2010; Wang et al., 2012). Additionally, in most splicing factor mutants, only small fractions of total transcripts are aberrantly processed (Jones et al., 2012; Perez-Santángelo et al., 2014; Feke et al., 2019; Li et al., 2019). Therefore, whether these small changes in levels of aberrantly spliced isoforms could lead to significantly decreased levels of functional proteins and thereby cause the observed circadian phenotypes remains unclear.

Here, we demonstrate that in *xct* mutants the circadian clock phenotype is genetically separable from altered levels of aberrantly spliced mRNA variants of core clock gene. We found that loss of *XCT* function accelerates the clock and causes aberrant splicing of five core clock genes (Supplemental Fig. 1; Fig. 4A). Surprisingly, the clock phenotype but not the splicing defects of the five clock genes in *xct-2* is suppressed by reduction of *PRP19A* or *PRP19B* function (Fig. 6). Similarly, the clock in *xct-1* runs slightly faster than *xct-2* but the splicing defects, including the aberrant splicing of *TOC1*, *TIC* and *CCA1*, are more severe in the latter (Fig. 2; Supplemental Fig. 5). Collectively, these data demonstrate that the short period phenotypes and altered levels of clock gene splicing variants are genetically separable in *xct* mutants.

How *XCT* regulates the pace of the circadian clock is still an outstanding question. Although we argue that changes in levels of aberrantly spliced clock mRNAs are not responsible for the accelerated clock in *xct* mutants, one possibility is that alterations in the overall kinetics of pre-mRNA splicing may cause circadian period phenotypes. Indeed, pharmacological perturbations of global transcription and translation efficiency can both lengthen the circadian period (de Melo et al., 2021; Uehara et al., 2022). Those results suggest that changes in the kinetics of RNA processing might cause the period phenotypes observed in plants mutated for some splicing factors.

Alternatively, *XCT* may control the circadian clock function independent of its role in pre-mRNA splicing. In Chlamydomonas, the XCT homolog XAP5 co-immunoprecipitates with RNA Pol II (Li et al., 2018), suggesting that XCT orthologs may possess transcriptional regulatory activities. In this paper, we show that NTC components physically and genetically interact with XCT to regulate circadian clock period (Table 1; Fig. 6A). Besides splicing, another well-characterized role of the NTC in gene expression is regulation of transcriptional elongation (Chanarat and Sträßer, 2013). Studies in both yeast and Arabidopsis have revealed that multiple NTC members, including PRP19, PRL1 and CDC5, physically associate with RNA Pol II and participate in transcript elongation (Chanarat et al., 2011; Zhang et al., 2013; Zhang et al., 2014). In addition, the NTC component SKIP interacts with Polymerase-Associated Factor 1 complex to regulate transcription in a splicing-independent manner (Li et al., 2019). Interestingly, a recent study showed that inhibition of transcriptional elongation by decreasing phosphorylation of the RNA Pol II C-terminal domain lengthens circadian period in Arabidopsis (Uehara et al., 2022). It is therefore possible that *XCT* affects circadian period by altering transcriptional elongation.

Yet another possibility is that *XCT* affects a cellular process independently of its role in gene expression. For example, PRP19 is known to facilitate DNA repair by its E3 ubiquitin ligase activity independent of its involvement in the spliceosomal complex (Maréchal et al., 2014; Idrissou and Maréchal, 2022). Indeed, there is increasing evidence that many RNA binding proteins directly participate in DNA double-strand break responses (Klaric et al., 2021). A recent study showed that the NTC-associated protein MOS4-ASSOCIATED COMPLEX SUBUNIT 5A (MAC5A) physically interacts with the 26S proteasome and regulates its activities in response to DNA damage (Meng et al., 2022). Intriguingly, we recently reported that loss of *XCT* function also disturbs the DNA damage response pathway (Kumimoto et al., 2021). Thus, it is possible that *XCT* works with *PRP19* in this pathway separately from its role in RNA processing. Future research is required to understand the nature of the relationship of between XCT and the NTC in the control of circadian clock function and other essential biological processes.

## Materials and Methods

### Plant materials and growth conditions

All *Arabidopsis thaliana* plants used in this study are Columbia-0 (Col-0) ecotype and contain a *CCR2*::*LUC* reporter for circadian clock assays. The *xct-1*, *xct-2*, *xct-2* p*XCT*::g*XCT-YFP-HA*, *prp19a-1*, *prp19b-1*, *prl1-2*, *cdc5-1* and *skip-1* genotypes have been previously described (Martin-Tryon and Harmer, 2008; Monaghan et al., 2009; Zhang et al., 2014; Wang et al., 2012). All the double mutants in this study were produced by crossing. Unless otherwise specified, seeds were surface sterilized with chlorine gas for 3 hours (50 ml 100% bleach + 3ml 1M HCl) and then plated on 1x Murashige and Skoog (MS) growth media containing 0.7% agar, pH 5.7. After 3d stratification in dark at 4°C, plates were transferred to 12-h light (cool white fluorescent bulbs, 55 µmol m^-2^ s^-1^) / 12-h dark cycles at 22°C for a variable number of days depending on the experiment.

### Immunoprecipitations and mass spectrometry

Plants were grown in 12-h light / 12-h dark cycles and harvested on day 10 at ZT3 or ZT17. Approximately 7.5g of vegetative tissue was flash frozen in liquid nitrogen and ground into fine powder with a mortar and pestle. Ground tissue was resuspended in nuclei enrichment buffer (50 mM Tris pH7.5, 400 mM sucrose, 2.5 mM MgCl_2_, 1 mM DTT, 1 mM PMSF, cOmplete Protease Inhibitor cocktail (Roche)), the nuclear pellet was collected, resuspended in IP buffer (100mM Tris pH7.5, 1mM EDTA, 75mM NaCl, 10% glycerol, 0.3% Triton X-100, 0.05% SDS, 10 mM MG-132, 1 mM PMSF, , cOmplete Protease Inhibitor cocktail (Roche)) and pelleted again. Nuclei were resuspended in IP buffer, disrupted via sonication, and adjusted to 1 mg/mL in IP buffer. 1 mg of extract was incubated with µMACS MicroBeads conjugated to an anti-HA monoclonal antibody (Miltenyi Biotec) and beads were then captured on µMACS M-colums. Beads were washed 3x with ice cold IP buffer and 1x with ice cold TE buffer (10 mM Tris-HCl pH 8.0, 0.1 mM EDTA). Proteins were eluted off the beads with elution buffer (Miltenyi Biotec). Proteins were resolved using SDS polyacrylamide gel electrophoresis and gels were sent to the Rutgers Biological Mass Spectrometry Facility, Robert Wood Johnson Medical School. Proteins were eluted and subjected to mass spectrometry as previously described (Wei et al., 2020).

### PacBio Iso-Seq and bioinformatic analysis

Arabidopsis seeds were surface sterilized in 70% ethanol prepared in 0.1% (v/v) Triton X-100 (Sigma) for 5 minutes and then in 100% Ethanol for 20 minutes. After air drying, seeds were plated on 1x MS growth media containing 0.7% agar. After cold stratification for 3 days, plates were transferred to constant light (55 µmol m^-2^ s^-1^) and temperature (22°C). Approximately 60 mg of whole seedlings of each genotype were collected every 2h over 24 hours on day 11 and pooled as indicated in Fig. 2A. Three biological replicates were collected for each genotype. Samples were flash-frozen in liquid nitrogen, and ground into fine powder using a beadbeater. Total RNA was extracted using an RNeasy Plant Mini Kit (Qiagen) followed by purification using the RNA Clean & Concentrator Kit (Zymo Research). Purified RNAs were quantified using a Nanodrop (ThermoFisher) and quality control was performed by an Agilent Bioanalyzer 2100. All samples had a 260/230 nm ratio higher than 1.9, 260/280nm ratio between 2 and 2.15, and RIN score higher than 8. PacBio Sequel II library preparation and RNA-sequencing (Iso-Seq) was performed by UC Davis DNA Technologies & Expression Analysis Core (https://dnatech.genomecenter.ucdavis.edu/pacbio-library-prep-sequencing/).

Raw reads generated by the PacBio Sequel II sequencer were imported into PacBio SMRT Link v 8.0 for Circular Consensus Sequence (CCS) calling and demultiplexing. Next, poly(A) tails and concatemers were removed using the refine command from isoseq3 package via Bioconda (v 3.10) with the option ‘--require-polya’. Then the fastq files containing full-length non-concatemer reads were mapped to Arabidopsis TAIR 10 genome assembly using minimap2 (v 2.17) with the parameter ‘-ax splice -t 30 -uf --secondary=no -C5’. For differential splicing analysis, counts of exonic regions and known/novel splice junctions were generated using QoRTs software package (v 1.3.6) (Hartley and Mullikin, 2015) with the parameter ‘--stranded -- singleEnded --stranded_fr_secondstrand --keepMultiMapped’. To adapt to the long-read data, the ‘maxPhredScore’ was set to 93 and the ‘maxReadLength’ was adjusted manually for each library. To increase the power of detecting novel splice junctions, the ’--minCount’ threshold was set to 5. The raw count data was then loaded to JunctionSeq R package (v 1.5.4) (Hartley and Mullikin, 2016) to determine differential usage of exons or splice junctions relative to the overall expression of the corresponding gene with a false discovery rate (FDR) of 0.05. Genes with at least one differentially spliced exon/junction were compared to the total circadian-clock-regulated genes (Romanowski et al., 2020) as described in Fig. 4.

### RNA extraction and RT-PCR

For time-course qRT-PCR analysis, Arabidopsis seedlings were entrained in 12-h light / 12-h dark cycles for 9 days before transferred to constant light and temperature (55 µmol m^-2^ s^-1^, 22°C). Starting from day 9, approximately 15-20 whole seedlings of each genotype were harvested every 4h over the 72-h period. For single time point gene expression and splicing analysis, 10-d-old seedlings grown in 12-h light / 12-h dark cycles at 22°C were harvested at estimated peak time of expression for each studied gene. Collected tissue was immediately frozen in liquid nitrogen and ground into fine powder using a beadbeater. Total RNA was isolated using TRIzol reagent (Invitrogen) and quantified with a Nanodrop (ThermoFisher). 200 ng of total RNA was used for cDNA synthesis with an oligo(dT)18 primer and SuperScriptIII (Invitrogen) reverse transcriptase. Diluted cDNAs were used as templates for qRT-PCR and semi-qRT-PCR reactions. The qRT-PCR was carried out as previously described (Martin-Tryon et al., 2007) using a Bio-Rad CFX96 thermocycler. Primers amplifying the aberrantly spliced or total transcripts were tested by standard curve and melt curve assays. Relative expression (ΔΔCq) values were normalized to the geometric mean of *PROTEIN PHOSPHATASE 2a* (*PP2A*) and *ISOPENTENYL DIPHOSPHATE ISOMERASE 2* (*IPP2*) expression levels. For semi-qRT-PCR, splice-junction specific primer pairs were designed to amplify regions flanking the aberrantly spliced introns of interest. The size and abundance of the resultant PCR products were then analyzed and quantified by LabChip GX bioanalyzer (PerkinElmer). The expression analyses represent two to three biological replicates. All primers used in this study are described in Supplemental Dataset 5.

### Circadian period analysis

After growing on MS plates under 12-h light / 12-h dark cycles for 6 days, Arabidopsis seedlings were sprayed with 3 mM D-luciferin (Biosynth) prepared in 0.01% (v/v) Triton X-100 (Sigma) and then transferred to a growth chamber with a constant 22°C and constant light provided by red and blue LED SnapLites (Quantum Devices, 35 µmol m^-2^ s^-1^ each) for imaging. Luciferase activity was detected using a cooled CCD camera (DU434-BV [Andor Technology]). Bioluminescence signals from the images were quantified using MetaMorph 7.7.1.0 software (Molecular Devices). Subsequent circadian period estimation and rhythmicity analysis were performed using Biological Rhythms Analysis Software System 3.0 (BRASS) by fitting the bioluminescence data to a cosine wave through Fourier Fast Transform-Non-Linear Least Squares (Plautz et al., 1997).

### Rosette size measurement

Seeds on MS plates were germinated in a growth chamber with 12-h light / 12-h dark cycles at a constant 22°C. 15-day-old seedlings were transferred to soil and grown in 16-h light (cool-white fluorescent bulbs, 75 µmol m^-2^ s^-1^) / 8-h dark long day condition at a constant 22°C. Rosette sizes of 39-day-old plants were determined by the greatest distance between rosette leaves using a digital caliper.

### Accession Numbers

All the *Arabidopsis thaliana* genes studied in this paper can be found under the following accession numbers:

*XCT*, AT2G21150

*PRP19A*, AT1G04510

*PRP19B*, AT2G33340

*PRL1*, AT4G15900

*SKIP*, AT1G77180

*CDC5*, AT1G09770

*LHY*, AT1G01060

*LNK2*, AT3G54500

*TIC*, AT3G22380

*PRR7*, AT5G02810

*TOC1*, AT5G61380

*SPPA*, AT1G73990

*IPP2*, AT3G02780

*PP2A*, AT1G69960

*CCA1*, AT2G46830

## Data Availability

All our MS data were deposited and available at Center for Computational Mass Spectrometry (CCMS, Dataset: MSV000090830). Our PacBio Iso-Seq data were deposited and available in NCBI Gene Expression Omnibus (GEO, Accession number: GSE220902).

## Acknowledgments

We thank the Arabidopsis Biological Resource Center for providing seeds. We thank Haiyan Zheng and the Biological Mass Spectrometry Facility of Robert Wood Johnson Medical School and Rutgers for mass spectrometry analysis. They are supported by NIH Shared Instrumentation Grant S10OD01640. We thank the DNA Technologies and Expression Analysis Core at the UC Davis Genome Center for performing PacBio Iso-Seq experiment. They are supported by NIH Shared Instrumentation Grant 1S10OD010786-01. We thank Julin Maloof for statistical advice, as well as members of the Harmer, Maloof, and Shabek labs for helpful discussions.

## Author Contributions

HZ, RWK and SLH designed the research; HZ, RWK, and SA performed the research; HZ, RWK, and SLH analyzed data; HZ and SLH wrote the paper.

## Supplemental Materials

**Supplemental Figure 1.** X***C***T **regulates plant development and the pace of the circadian clock in *Arabidopsis thaliana.*** A, Morphology of 52-day-old Col-0, *xct-1*, *xct-2* mutant plants and *xct-2* complemented with p*XCT*::g*XCT-YFP-HA.* Bar = 10 mm. B, Circadian periods of genotypes described in (A). The lines in the boxplot represent the 75% quartile, median and 25% quartile of the data, respectively. Statistical significance was determined using linear regression model with genotype as a fixed effect and is shown in lower case letters (Tukey’s multiple comparison test, *P* < 0.05).

**Supplemental Figure 2.** X***C***T **and *PRL1* control the pace of the circadian clock in an opposite manner.** A, Morphology of 21-day-old Col-0, *xct-1*, *xct-2* and *prl1-2* mutant Arabidopsis seedlings. Bar = 1 mm. B, Circadian clock periods of plant genotypes described in (A). The lines in the boxplot represent the 75% quartile, median and 25% quartile of the data, respectively. Statistical significance was determined using linear regression model with genotype as a fixed effect and is shown in lower case letters (Tukey’s multiple comparison test, *P* < 0.05).

**Supplemental Figure 3.** p***r***l1***-2* mutant mainly induces intron retention defects in Arabidopsis.** Frequency of different classes of 5’ and 3’ splice sites of down-(teal) or up-regulated (orange) junctions in *prl1-2* compared to wild type. Known or novel splice sites were classified based on TAIR10 genome annotation.

**Supplemental Figure 4.** x***c***t***-1* mutant displays reduced fidelity of 3’ splice site selection during pre-mRNA splicing.** A, Distribution of the distances between each mis-selected novel 3’ splice site and its corresponding canonical 3’ splice site in *xct-1*. B - C, Pictograms showing the frequency of nucleotides in the 23-mers sequences flanking the 3’ splice sites in splice junctions that are down-regulated or up-regulated in *xct-1*.

**Supplemental Figure 5.** T**h**e **splicing defects of core clock genes are more severe in *xct-2* than *xct-1*.** A - D, Sashimi plots showing PacBio Iso-Seq reads mapped to *CCA1* (A)*, LNK2* (B)*, TOC1* (C) and *TIC* (D) in Col-0 (teal)*, xct-1* (yellow) and *xct-2* (orange). The red rectangles highlight the aberrantly spliced exon-exon junctions that are examined by semi-qRT-PCR in (E) - (H). E - H, Relative abundance of aberrantly-spliced to fully-spliced isoforms of *CCA1* (E)*, LNK2* (F)*, TOC1* (G) and *TIC* (H). Differentially spliced isoforms were amplified by semi-qRT-PCR using primers flanking the examined regions, followed by separation and quantification using a Lab Chip GX bioanalyzer. Statistical significance was determined using linear regression models with genotype as a fixed effect and is shown in lower case letters (Tukey’s multiple comparison test, *P* < 0.05).

**Supplemental Figure 6.** D**a**wn**-phased genes are significantly enriched among genes mis-spliced in *xct-1.*** A - B, Phases of peak expression of all the circadian-clock-regulated genes (Romanowski et al., 2020) that are detected (A) or aberrantly spliced (B) in *xct-1*. The white and gray backgrounds represent the subjective day and subjective night, respectively. C, Distribution of *P*-values from Fisher’s exact test calculating whether the ratio of the number of mis-spliced genes in (B) to all clock-regulated genes in (A) is significantly higher than expected by chance for each 2-hour interval.

**Supplemental Figure 7.** Loss of *XCT* function does not affect overall abundance of transcripts of core-circadian-clock genes that are aberrantly spliced in *xct* mutants. A - F, Normalized expression of the total transcript levels of *LHY* (A)*, LNK2* (B), *PRR7* (C) and *TOC1* (D) in Col-0*, xct-2* and *xct-2 XCT*. Samples were collected every four hours over a 72-h window. Expression levels were examined by qRT-PCR using primers that detect abundance of all transcripts and normalized to *PP2*A and *IPP2.* Data points represent mean ± se from three independent biological replicates. For each isoform in each biological replicate, the normalized expression levels were relative to the highest expression level in Col-0 across all time points. Teal lines, wild type Col-0; orange lines, *xct-2* muants; purple lines, *xct-2* complemented with p*XCT*::g*XCT-YFP-HA*. Black background, dark period; white and gray background, light period during subjective day and night, respectively.

**Supplemental Figure 8.** M**u**tation **of Arabidopsis NTC members lengthens circadian period.** A - C, Circadian periods of Col-0 and NTC mutant plants, including (A) *prp19a-1*, *prp19b-1*, *prp19a-1 prp19b-1*, (B) *cdc5-1*, and (C) *skip-1*. The lines in the boxplot represent the 75% quartile, median and 25% quartile of the data, respectively. Statistical significance was determined using linear regression model with genotype as a fixed effect and is shown in lower case letters (Tukey’s multiple comparison test, *P* < 0.05).

**Supplemental Figure 9.** P*R*P19A (*MAC3A*) and *PRP19B* (*MAC3B*) transcript levels are highly reduced in *prp19a-1* and *prp19b-1* mutant backgrounds, respectively. A - B, Normalized expression of (A) *PRP19A* and (B) *PRP19B* in Col-0, *xct-2*, *prp19a-1*, *prp19b-1*, *prp19a-1 prp19b-1*, *prp19a-1 xct-2* and *prp19b-1 xct-2*. Expression levels were examined by qRT-PCR and normalized to *PP2*A and *IPP2.* Data points represent mean ± se from two independent biological replicates. Samples were collected at ZT0. Statistical significance was determined using linear regression model with genotype as a fixed effect and is shown in lower case letters (Tukey’s multiple comparison test, *P* < 0.05).

**Supplemental Figure 10.** L**o**ss **of *PRP19* function suppresses the *xct-2* short circadian period phenotype.** An independent replicate of the experiment in Figure 6A showing the circadian periods of Col-0, *xct-2*, *prp19a-1*, *prp19b-1*, *prp19a-1 prp19b-1*, *prp19a-1 xct-2* and *prp19b-1 xct-2* plants.

**Supplemental Figure 11.** P***R***P19 **and *XCT* additively regulate Arabidopsis rosette size.** A, Morphology of 39-day-old Col-0, *xct-2*, *prp19a-1*, *prp19b-1*, *prp19a-1 prp19b-1*, *prp19b-1 xct-2* and *prp19a-1 xct-2* plants. Bar = 10 mm. B, Rosette diameter of 36-day-old plants grown under long-day conditions at 22°C (*n* = 5-11). Data are shown as mean ± se. Statistical significance was determined using linear regression model with genotype as a fixed effect and is shown in lower case letters (Tukey’s multiple comparison test, *P* < 0.05).

**Supplemental Figure 12. Levels of total transcripts of *XCT*-targeted core clock genes are not altered in *prp19a-1 xct-2* or *prp19b-1 xct-2* compared to *xct-2.*** A - F, Normalized expression of total transcript levels (A, C, E, F) or specific normally spliced isoforms (B, D) of *LHY, LNK2, TIC, PRR7* and *TOC1* in Col-0, *xct-2*, *prp19a-1*, *prp19b-1*, *prp19a-1 prp19b-1*, *prp19a-1 xct-2* and *prp19b-1 xct-2*. Expression levels were examined by qRT-PCR using primers that detect abundance of all transcripts or normally spliced isoforms and normalized to *PP2*A and *IPP2.* Data points represent mean ± se from two independent biological replicates. Samples were collected at the estimated peak expression time for each gene. Statistical significance was determined using linear regression model with genotype as a fixed effect and is shown in lower case letters (Tukey’s multiple comparison test, *P* < 0.05).

Supplemental Dataset 1. A list of proteins co-purified with XCT-IP detected by MS.

Supplemental Dataset 2. Summary of full-length reads obtained from each PacBio Iso-Seq library.

Supplemental Dataset 3. Lists of significant differential splicing events identified by JunctionSeq.

Supplemental Dataset 4. Lists of total and aberrantly spliced circadian-clock-regulated genes described in this study.

Supplemental Dataset 5. A list of primers used in this study.

